# BQsupports: systematic assessment of the support and novelty of new biomedical associations

**DOI:** 10.1101/2023.03.09.531915

**Authors:** Adrià Fernández-Torras, Martina Locatelli, Martino Bertoni, Patrick Aloy

**Affiliations:** Institute for Research in Biomedicine (IRB Barcelona), The Barcelona Institute of Science and Technology, Barcelona, Catalonia, Spain; Institució Catalana de Recerca i Estudis Avançats (ICREA), Barcelona, Catalonia, Spain

## Abstract

Living a Big Data era in Biomedicine, there is an unmet need to systematically assess experimental observations in the context of available information. This assessment would offer a means for an unbiased validation of the results and provide an initial estimate of the potential novelty of the findings. Here we present BQsupports, a web-based tool built upon the Bioteque biomedical descriptors that systematically analyzes and quantifies the current support to a given set of observations. The tool relies on over 1,000 distinct types of biomedical descriptors, covering over 11 different biological and chemical entities, including genes, cell lines, diseases and small molecules. By exploring hundreds of descriptors, BQsupports provide support scores for each observation across a wide variety of biomedical contexts. These scores are then aggregated to summarize the biomedical support of the assessed dataset as a whole. Finally, the BQsupports also suggests predictive features of the given dataset, which can be exploited in downstream machine learning applications.

**Availability and implementation:** The web application and underlying data are available online (https://bqsuppports.irbbarcelona.org).

## Introduction

Since the popularization of high-throughput experiments to obtain an unbiased and more comprehensive description of biology, many initiatives have massively gathered data from biological systems^1^. Initial efforts relied on model organism screenings to uncover protein-protein interactions^2^ and gene co-expression patterns^3^. Concurrently, drug repositories began to annotate bioactivity data for hundreds of drugs^4-6^. With the consolidation of large cell line panels, traditional OMICS started to gather all sorts of biological descriptors, from mutations in the genome to protein abundances^7,8^. The next generations incorporated global biological responses to small molecules and genetic perturbations^9-11^. Eventually, the accumulation of genomic data enabled the statistical exploration of gene-phenotype associations, leading to the extensive identification of disease-associated genes^12,13^.

All these initiatives are populating biomedical repositories with hundreds of datasets, many of which are part of monumental efforts that still keep providing new releases to date (e.g.^14-17^). Many of these initiatives are gigantic efforts that take decades to complete and, while first-in-class datasets often offer a wealth of new biological findings, it is important to assess the novelty of subsequent releases to determine the optimal screening strategies. Thus, it is paramount to have the means to contextualize new data in light of current biomedical knowledge. Indeed, it is a common practice to inspect experimental results based on existing data, as it helps to validate new methodologies and the results, thereby, gaining confidence in the provided insights. Unfortunately, no standard exists for this analysis, which inevitably hampers the unbiased comparison with existing resources. Besides, these assessments usually use previous releases or analogous datasets as a reference, missing whether similar relationships have been found in orthogonal studies.

Here, we present *Bioteque Supports* (*BQsupports)*, a web tool to systematically quantify the support of novel biological associations based on the current biomedical knowledge pre-encoded in the Bioteque^18^. BQsupports assesses the similarity of each pair of biomedical entities given by the user within a collection of diverse Bioteque descriptors, providing support scores across various biomedical contexts. Additionally, by identifying the descriptors that better explain each novel type of association, BQsupports suggests biomedical features that can be used to predict relationships between entities that the screens might have missed. Overall, by exploring hundreds of descriptors, BQsupports provide support scores for novel experimental observations across a wide variety of biomedical contexts.

## BQsupports description

BQsupports provides biomedical support scores between pairs of biological entities given by the user. This support derives from biomedical knowledge descriptors (i.e. *embeddings*) gathered in the Bioteque resource^18^, which can assess links between 11 different types of biomedical entities, namely: genes/proteins (GEN), cell lines (CLL), tissues (TIS), small molecule compounds (CPD), diseases (DIS), pharmacological classes (PHC), chemical entities (CHE), pathways (PWY), cellular components (CMP), protein domains (DOM) and molecular functions (MFN). Experimentally determined relationships between these biomedical entities (i.e. *nodes*) are then collected, harmonized and embedded in the form of context-dependent biomedical descriptors (i.e. *metapaths*). We have pre-computed over 1,000 metapath embeddings, so that a given biological entity can be described in many different ways, depending on its context. For instance, we have different descriptors for a protein accounting for its pattern of interactions, the cells or tissues where it is expressed, the diseases where it is involved or the drugs targeting it. Overall, nodes that are close in a given metapath space show similar biomedical properties in that context.

Given a set of node pairs covered by the Bioteque, BQsupports automatically identifies contextual biomedical descriptors, or metapaths, potentially related to the input data. Then, within each context space, it measures the cosine distance between nodes in each pair and provides a support score for each pair. These individual, context-related, scores are further aggregated to obtain a single support estimate for the entire dataset. Moreover, the tool automatically runs a network permutation protocol to (i) derive the expected support of the dataset and (ii) detect entity pairs that are significantly close (supported) in a particular biomedical context, which are quantified by an enrichment score. Additionally, BQsupports identifies metapaths able to distinguish the dataset associations from random permutations, thus providing means to prospectively predict associations that might be missing in the input dataset. All the analyses and results for each input data can be downloaded in the form of different tables and they are summarized in a canvas picture. The entire pipeline is detailed in the Supplementary Methods section.

As an illustrative example, Figure 1 shows the BQsupport results for the Bioplex-III protein-protein interaction (PPI) network19. The heatmap on the left shows the level of support that different metapaths (rows) give to each PPI in the dataset (columns). When considering all the biomedical context descriptors together (last row), we see that over 73% of the interactions are supported by current knowledge (quantile rank ≤ 0.05). In other words, 27% (19.1k) of the PPIs identified in the Bioplex-III network can be considered potentially novel, according to our resource. The expected support suggests that the interactions in this dataset are more backed up by other observations than expected by chance, especially within the stronger (redder) support range. The TOP metapath ranking shows that most of this support comes from descriptors based on previously known PPIs (GEN-ppi-GEN), followed by descriptors finding the provided interactions in similar cell compartments (GEN-has-CMP) and pathways (GEN-ass-PWY). Not surprisingly, it is also these three metapaths the ones harboring a higher potential for predicting PPIs that the current versions of Bioplex might have missed.

**Figure 1.**
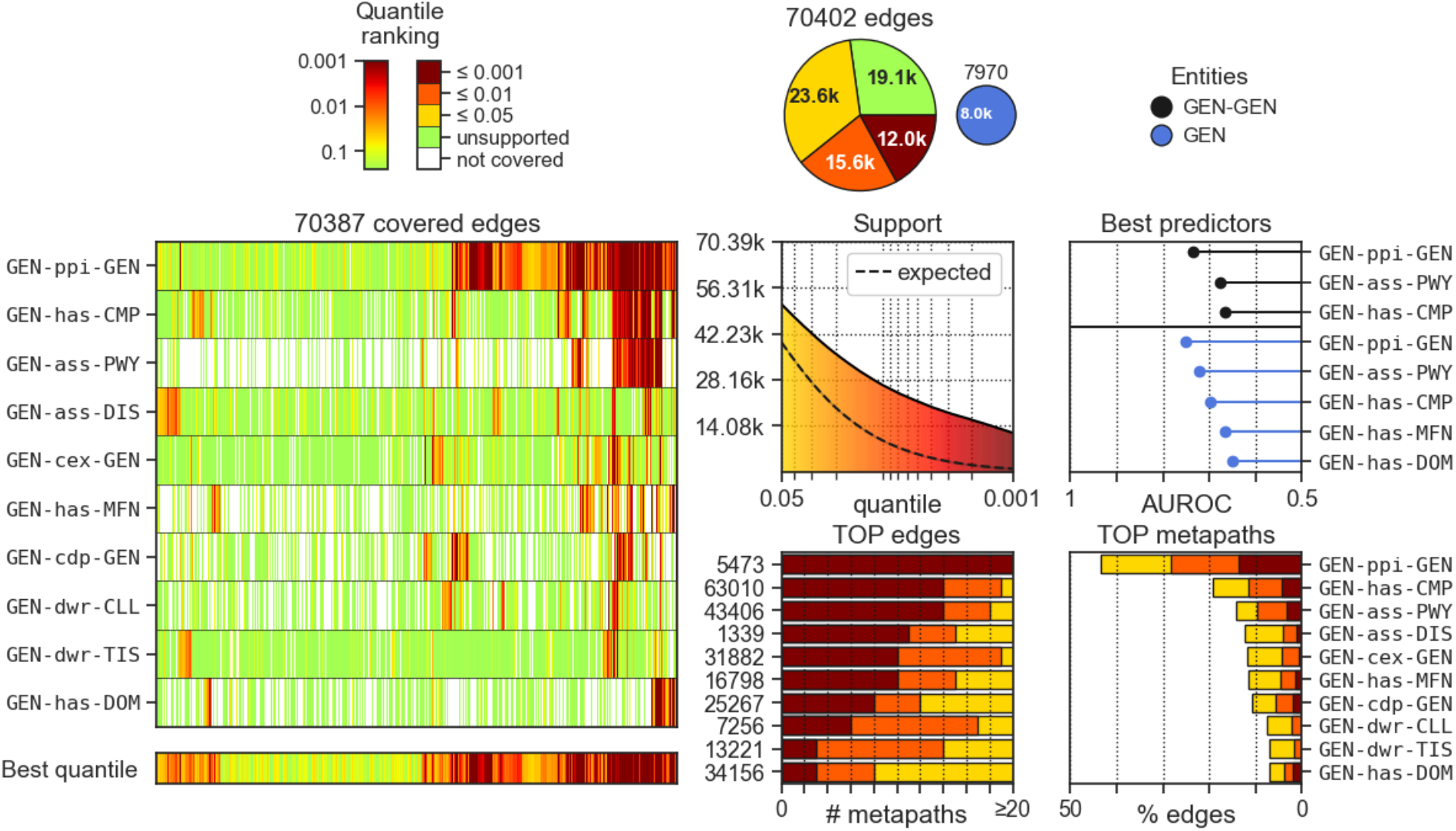
BQsupports analytical canvas for the Bioplex protein-protein interaction data. On the left, we show the quantile ranking (support score) for all the input relationships (y-axis) covered by the top 10 most supportive metapaths (x-axis). The lower (redder) the quantile rank, the higher the support score. On the right, we stratify the support scores according to different cutoffs, and summarize the results in a pie chart, along with the coverage of the input dataset (minor pie chart). Below, we quantify the support of the input dataset across different quantiles, showing the ‘expected support’ achieved by permuted networks (i.e. random expectation). Next to it, we rank the most predictive metapaths, quantified as the Area Under the ROC (AUROC). We perform this analysis for the specific pairs provided in the input dataset (black) and for entity types pairs sharing a similar association profile (Supplementary Methods). Finally, we show the top most supported input edges and the metapaths that most support the dataset at the bottom right corner.

## Concluding remarks

BQsupports offers a means to systematically assess the novelty of new experimental data, providing support scores for each observation in a given dataset while identifying features suitable for downstream predictive tasks. BQsupports is available as a web-based tool, where the user is only asked to (i) provide the dataset associations with proper identifiers, (ii) specify the types of biomedical entities and (iii) optionally tune some parameters (e.g., the number of permuted networks). With default parameters, the whole BQsupports pipeline takes between 1h to 2h to complete, depending on the number of relationships provided (Supplementary methods). At the end of the process, BQSupports returns a canvas figure summarizing the results (Figure 1 is the direct output of the pipeline ran on Bioplex-III) and three table files covering the quantile ranking score for each association-metapath combination, a digested summary counts for each descriptor, and the estimated performance of these metapaths in downstream predictive tasks. Indeed, the provided files allow the user to easily recompute most of the presented analyses according to custom needs (e.g., limiting the support score to a particular set of biomedical contexts or requiring a minimum enrichment score). Also note that all the metapath descriptors can be downloaded from the main Bioteque page (https://bioteque.irbbarcelona.org).

## Acknowledgements

P.A. acknowledges the support of the Generalitat de Catalunya (RIS3CAT Emergents CECH: 001-P-001682 and VEIS: 001-P-001647; and 2021 SGR 00876), the Spanish Ministerio de Ciencia, Innovación y Universidades (PID2020-119535RB-I00), the Instituto de Salud Carlos III (IMPaCT-Data), and the European Commission (RiPCoN: 101003633). AF-T is a recipient of an FPI fellowship (BES-2017-083053). We also acknowledge institutional funding from the Spanish Ministry of Science and Innovation through the Centres of Excellence Severo Ochoa Award, and from the CERCA Programme / Generalitat de Catalunya.

## Author Contributions Statement

A.F.-T. and P.A. designed the study and wrote the manuscript. A.F.-T. implemented the entire computational strategy and analysis. A.F.-T, M.L. and M.B. implemented the web resource. All authors analyzed the results, and read and approved the manuscript.

## Supplementary Data

### BiotequeSupport coverage

BQsupports fetches biomedical descriptors from the Bioteque resource^18^. At the date of publication, it covers 11 different entities (nodes), allowing for the screening of 11 homogeneous and 110 bipartite entity-entity combinations (edges). Each entity type has its identifier vocabulary, enabling a harmonized data integration. The available entities, vocabularies, and metapath descriptors are specified in the web resource (https://bioteque.irbbarcelona.org).

### Scalability

We tested the tool with tens of different datasets and confirmed that it scales well with hundreds of thousand edges. However, we capped the maximum number of edges to 1M to control the computational resources. With the default number of permutations (20) to estimate the expected level of support, the entire pipeline takes between 1h to 2h to run, depending on the entity types and number of edges (Figure S1, left). However, we observed that the number of permuted networks significantly impacts the computational time. For example, in the Bioplex-III network, moving from 20 to 100 random permutations added two hours of extra computation, while moving from 100 to 1000 added 18h more (Figure S1, right).

**Figure S1.**
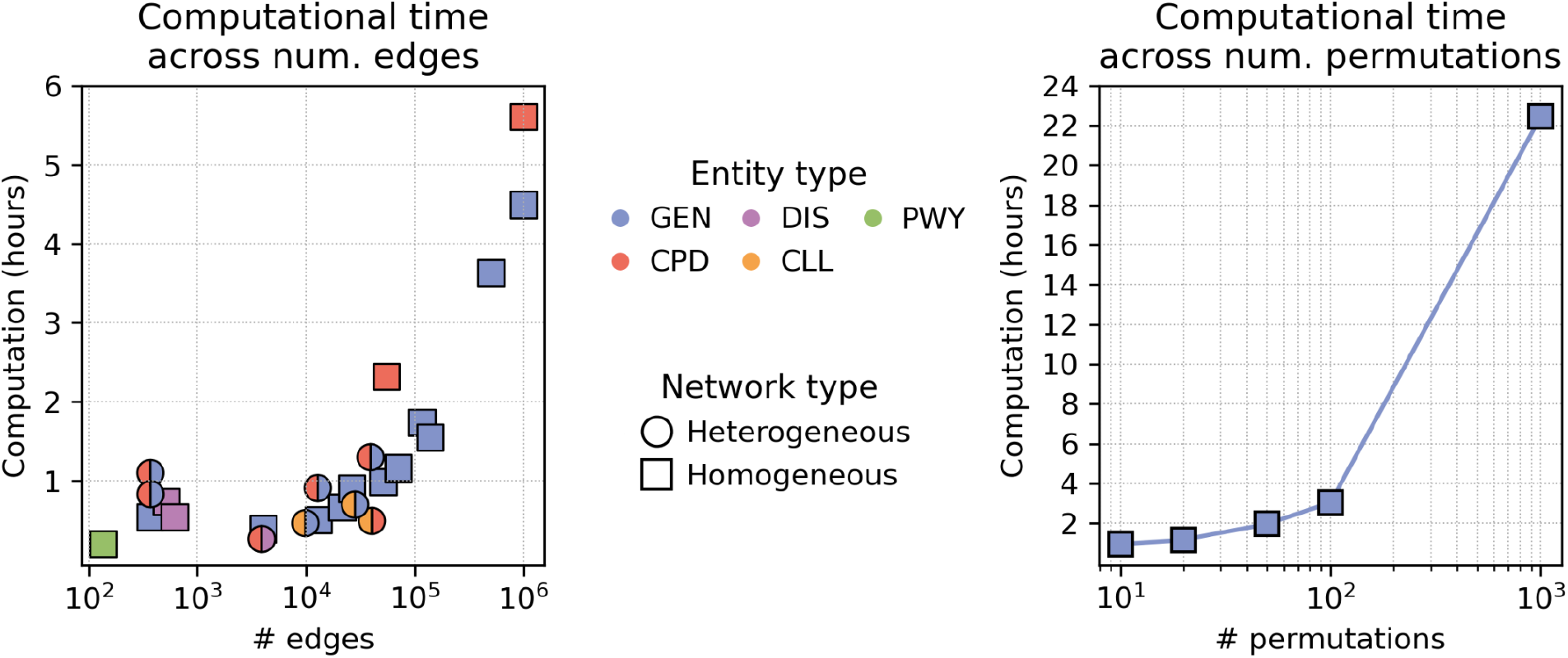
The computational time of the BQsupports pipeline. Left) Computational time in hours (y-axis) taken to run the full pipeline for different networks of varying size (x-axis) and type (shape and color). Right) Computational time in hours (y-axis) taken to run the BQsupports pipeline on the Bioplex-III network using 10, 20, 50, 100, and 1000 network permutations (x-axis).

### Methodology

#### Data input and user interface

BQsupports accepts pairs of associated nodes (networks) as input data. Users can provide the data either explicitly to the web or by uploading an edge file. Next, the user has to specify the type of entities provided. When providing homogeneous networks, it is possible to specify whether the associations are undirected (e.g. protein-protein interactions) or directed (kinase-substrate interactions). This will affect the network permutation process (e.g. in a directed kinase-substrate network, random permutations will always produce kinase-substrate pairs). Users can also vary the number of permuted networks from 10 to 1000. By default, the tool uses 20 network permutations, allowing a p-value resolution of 0.05. Note that the number of permuted networks directly impacts the enrichment score, where statistical power increases proportionally to the number of permutations (i.e. enrichment scores tend to be more significant and accurate). However, increasing the number of permuted networks will also affect the computational time of the pipeline (Figure S1, right). Lastly, it is important to note that BQsupports will skip repeated edges, as they can artificially increase the support of the dataset. If the provided network is undirected, the pipeline will sort each edge before removing duplicates. At the end of the process, the final network is provided to the user.

#### Calculation of support scores

The pipeline starts by listing all the metapaths connecting the entities specified by the user. It considers metapaths of any length available in the Bioteque resource, except for GEN-GEN associations, which are limited to L1 metapaths. BQsupports omits metapaths covering less than 10% of the data.

Once the metapath universe is defined, it computes cosine distances between each provided association in each metapath space and ranks them according to the metapath distance distribution. To obtain these rankings efficiently, only the top 25% closest neighbors (first quartile) for each node are retrieved using FAISS^20^. Accordingly, the node found in the first quartile sets the maximum ranking distance in the metapath. Next, BQsupports transforms rankings into quantiles by dividing them by the number of nodes in the embedding space. Finally, as this process generates two quantiles (i.e. one ranking for each node), it derives an edge-level quantile by keeping the geometric mean of the pair (i.e. the normalized co-rank). This process is repeated independently for each metapath-dataset descriptor in the pre-selected universe.

#### Calculation of random permutations and enrichment scores

To generate random permutations of the data, we perform *n* random swaps of the network using the BiRewire Bioconductor package^21^, where *n* is fixed to be ten times the number of edges in the dataset. Then, quantile ranking scores are calculated independently for each network permutation following the pipeline described in the previous section.

Enrichment scores are computed for each metapath and support scores. More specifically, given a metapath-source descriptor space and a quantile cutoff (tested range between 1 and 0.001), the pipeline first annotates the number of associations in the given dataset that score lower than the given quantile cutoff. Then, it obtains a Fold Change (FC) by dividing this number by the median number of associations obtained from the random permutations. Additionally, it derives an empirical p-value by counting the proportion of permuted networks with equal or more associations than the original dataset. Notice that the resolution of this p-value will depend on the number of permuted networks (e.g. given 20 random permutations, the lowest computable p-value is *< 0*.*05*).

#### Identifying the best metapaths to complete missing node associations

To suggest the prediction potential of the different metapaths, the tool evaluates the capacity of metapath descriptors to distinguish the dataset associations from random permutations by ranking all the associations according to their cosine similarities. In those embedding spaces where ‘real’ edges (i.e. those given by the user) are up-ranked before random permutations, we can assume that the space preserves the structure of the dataset. Thus, the descriptors of this metapath space are likely to embed useful information to predict such type of relationships between the nodes. This is quantified by computing the Area Under the Receiver Operating Characteristic (AUROC) curve between the user edges and 10 random permutations. To prevent an association from being counted as a positive and negative instance simultaneously, the pipeline generates new random permuted networks without allowing them to overlap with the data provided by the user. At the end of the process, BQsupports provides the AUROC average across the 10 permutations together with the universe of each metapath and the covered portion of the input dataset. Note that the covered data represent the applicability domain of the computed AUROC.

Additionally, the pipeline also looks for metapath descriptors that, while not directly preserving the associations provided by the user, retain the neighborhood similarity of the nodes. In other words, BQsupports first identifies pairs of nodes with similar interactions than the supplied by the user, and then tries to find metapath descriptors that recapitulate these pairs. To identify these common interactors, the pipeline builds a similarity network by linking the input nodes with other nodes of the same entity type that share a significant number of associations. This similarity network is created by (i) representing each node with a binary vector annotating their interactions (i.e. the adjacency matrix), (ii) calculating term frequency-inverse document frequency (TF-IDF) values between the vectors and (iii) keeping the top 3 neighbors with highest TF-IDF similarity for each node. As a result, a new homogenous network is obtained, whose edges capture the most similar nodes from the network provided by the user. Next, the pipeline lists a new metapath universe for this network and computes the recapitulation (AUROC) scores of these metapaths, using the same approach described above. As a result, BQsupports identifies metapaths that keep the similarities between the nodes. Note that metapaths identified in this analysis may be different from those found to yield the highest prediction potential, even for homogeneous networks. For instance, in the example provided in Figure 1, the suggested entity predictors include two metapaths, namely GEN-has-MFN and GEN-has-DOM, that were not covered by the previous analysis. This would indicate that genes having similar molecular functions (GEN-has-MFN) or sharing protein domains (GEN-has-DOM) may not necessarily interact with each other but tend to interact with the same proteins, to some extent.

#### Output canvas generation

1. *Heatmap*. To generate the heatmap matrix, BQsupports first aggregates the scores by keeping the best quantile ranking among the sources belonging to the same metapath, obtaining a unique score per metapath. Then, it ranks metapaths according to the number of interactions they support with a quantile lower than 0.05, selecting the top 10 for the heatmap. Additionally, it provides the best quantile across all the screened metapaths in the last row. Associations not covered by a given metapath are left blank. Note that quantile scores are capped at 0.25 (1st quartile), as higher quantiles are ignored by the pipeline.
2. *Pie charts*. Support scores for each dataset association are aggregated by selecting the best score across metapaths. Next, they are stratified into four groups according to their quantile: ≤0.001, ≤0.01, ≤0.05, and *unsupported* (quantile > 0.05). The pie chart reports the counts of each group, together with those not covered by the resource (if any). Additionally, a minor pie chart depicts the fraction of nodes covered for each entity, colored according to the color code used in the Bioteque resource.
3. *Dataset support*. The total number of supported associations in the dataset is reported across the range of significant quantile rankings (from 0.05 to 0.001). Additionally, BQsupports annotates the mean and standard deviation achieved with permuted networks (dashed line), providing the expected supportiveness according to the dataset’s applicability domain (universe).
4. *Edge and metapath ranking*. The top 10 most supported edges and supportive metapaths are ranked according to the number of metapaths (or edges) with a quantile lower than 0.05. BQsupports uses the index provided in the original network (starting the count from 1) to label the edges in the plot (shown on the y-axis).
5. *Best metapath predictors*. The canvas shows the top 3 metapath for each tested network (i.e. the one provided by the user and each entity-entity similarity network generated by BQsupports). In this case, the reported scores are not aggregated by metapath and they correspond to a specific metapath-source combination. Furthermore, the tool only shows significant and relevant metapath-source combinations, those whose average AUROC value (after subtracting their standard deviation) is higher than 0.6 and cover at least 20% of the dataset.
6. Dataset results. In addition to the summary canvas, the pipeline outputs the results in different (.tsv) files. A first file provides, for each association-metapath-source triplet combination, (i) the quantile ranking score, (ii) cosine distance (with its corresponding Z-score transformation), and (iii) the inferred enrichment score (with its corresponding p-value) as detailed in the ‘*Random permuted networks and enrichment score inference’* section. A second file summarizes the number of edge counts supported by each metapath-dataset across different support scores. Lastly, a third file provides the recapitulation score (AUROCs) for each metapath assessed as a potential predictor, together with the coverage of the dataset and other practical information (e.g. metapath universe size).

